# Contingency degradation overwrites initial learning and depends on lateral orbitofrontal cortex

**DOI:** 10.64898/2026.05.18.726131

**Authors:** Maedeh Mahmoudi, Joanne M. Gladding, Michael D. Kendig, Alessandro Castorina, Karly M. Turner, Octavia Soegyono, Laura A. Bradfield

## Abstract

Relapse after treatment for various mental health disorders has been linked to tendency for reductions in responding to increase over time or following re-exposure to motivating stimuli. Here we show that, in rats, responding reduced through non-contingent outcome delivery does not recover in these ways, and that this learning depends on an intact lateral orbitofrontal cortex. These findings suggest that contingency degradation overwrites original learning which may support the development of relapse-resistant behavioural interventions.

---

Responding that has been reduced through experience is known to recover over time or following re-exposure to a biologically relevant outcome. First described by Pavlov in 1927^1^, these phenomena – spontaneous recovery and reinstatement – have since been extensively documented across species, most commonly^2,3^ (but not exclusively^4–6^) following extinction, during which a cue or action is no longer paired with its motivating outcome. Their discovery led to the widely accepted view that extinction temporarily suppresses rather than erases prior learning^3,7^, and that memory retrieval is dynamic and context-dependent^8^. Clinically, these phenomena are thought to contribute to relapse following behavioural treatments for anxiety, addiction and related disorders^9,10^. Identifying a procedure that reduces responding without recovery would therefore have broad theoretical and clinical significance.

A recent study revealed that contingency degradation might weaken response-outcome learning^11^, revealing it as a potential candidate for such a procedure. The present study therefore tested whether degrading the contingency between actions and outcomes produces a long-lasting reduction in responding, then examined the content and neural basis of this learning. Specifically, we assessed the behavioural properties of degradation using choice tests and outcome-selective reinstatement, then tested its dependence on lateral orbitofrontal cortex (lOFC) using chemogenetic inactivation.

We first trained male and female rats to press a right lever for pellets and a left lever for sucrose, or the opposite arrangement, counterbalanced (Fig 1A). Once press rates reached sufficient levels (Fig. 1B, main effect of day, F(1.557, 63.85) = 46.318, p < .001, no main effects of lever or time-of-test groups, and no interactions with either factor, Fs < 1.02), one lever press-outcome contingency was degraded through unearned, unsignalled deliveries of its outcome. Importantly, to avoid confounding degradation with extinction, both levers also continued to earn their respective outcomes at the same rate during the degradation phase. For example, if pellets were degraded, rats would earn pellets at similar rates whether they pressed the pellet lever or not, but to earn sucrose they still had to press the sucrose lever (the opposite was true if sucrose was degraded). This reduced responding on the degraded lever relative to the nondegraded lever (Figs. 1C-D), supported by a main effect of degradation, F(1,41) = 4.87, p = 0.033, and no main effects or interactions with time-of-test group (immediate vs. delay), Fs < 1.

**Figure 1.**
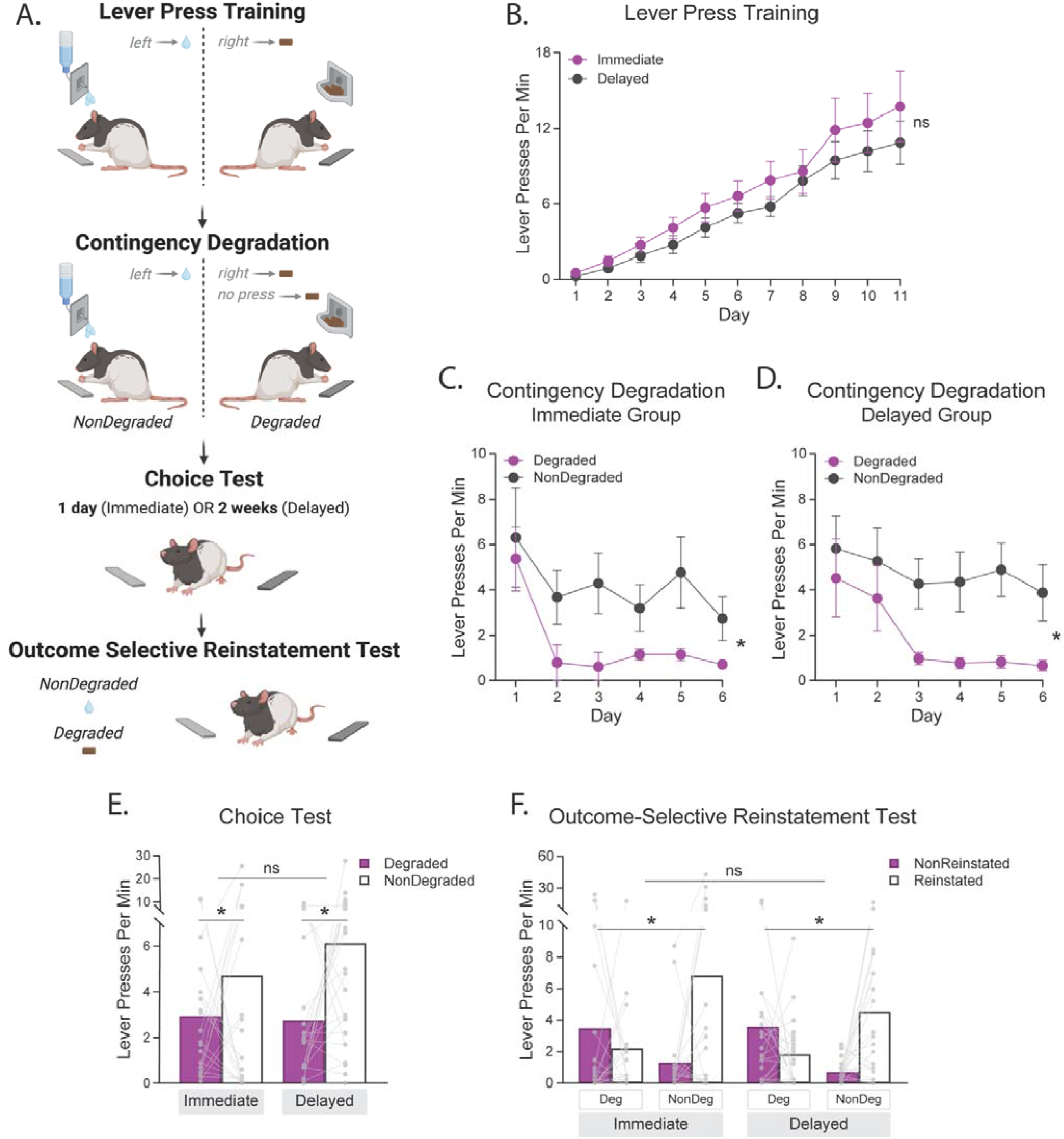
Contingency degradation does not spontaneously recover or reinstate. A) Design for Experiment 1. Top panel: rats were first trained to press left and right levers for distinct outcomes (e.g., left–sucrose, right–pellets). Upper middle panel: during contingency degradation, both levers continued to earn their outcomes, but one outcome (e.g., pellets) was also delivered non-contingently to degrade its lever–outcome relationship. Lower middle panel: rats then received a choice test either 1 day (immediate) or 2 weeks (delayed) later, conducted in extinction. Bottom panel: outcome-selective reinstatement was assessed with both levers available; each outcome was delivered twice across four trials (pellet, sucrose, sucrose, pellet) and post-delivery reinstatement was measured on each lever. B) Lever presses per min (± SEM) during lever press training in Experiment 1. C) Lever presses per min (± SEM) during contingency degradation for immediate group and D) for delayed group in Experiment 1. E) Individual data plots and mean lever presses during choice test, F) reinstatement test following the delivery of Degraded and NonDegraded outcomes. * p < 0.05.

Testing consisted of a ten-minute choice test in which both levers were presented and no outcomes earned. If degradation is resistant to spontaneous recovery, testing at each time point should reveal reduced responding on the degraded relative to the nondegraded lever, and levels of responding on the degraded lever should be equivalent across tests. As shown in Figure 1E, this was the observed result. Statistically, there was a main effect of degradation, F(1,41) = 4.69, p = 0.036, with no main effect of test delay, and test delay x degradation interaction, Fs < 1. This result confirmed that responding reduced by contingency degradation does not spontaneously recover.

We then conducted additional reinstatement tests at each time point, in which outcomes were unexpectedly delivered and the effect on responding measured. Like spontaneous recovery, reinstatement has also been extensively replicated across species and paradigms^12–14^, most commonly after extinction. When tested after a delay, reinstatement and spontaneous recovery have been shown to summate, producing an even greater increase in responding than either phenomenon alone^15^. Therefore, we theorised that if degradation produced a particularly robust form of response reduction, it would fail to restore responding even after a 2-week delay. The outcome-specific version of reinstatement used here additionally allowed us to probe the content of degradation learning. That is, pellet delivery will typically selectively increase responding on the pellet-associated lever, and likewise for sucrose^14,16^; we therefore asked whether this would still occur after the lever-outcome relationship had been degraded.

This was the pattern (i.e. Reinstated > NonReinstated, Fig. 1F) observed following deliveries of the nondegraded outcome at both time points. Remarkably, however, not only did deliveries of the degraded outcome fail to produce reinstatement on the degraded/reinstated lever, but it appeared to reverse the allocation of responding (NonReinstated > Reinstated, Fig. 1F) indicating that deliveries of this outcome *suppressed* responding on the degraded lever. These findings indicate that whereas the relationship between the nondegraded lever and its outcome was intact and excitatory at both time points, the relationship between the degraded lever and its outcome may have become partially inhibitory and remained so over time. Statistical support for these observations was provided by a main effect of reinstatement, F(1,41) = 16.49, p < .001, that interacted with degradation (reinstatement x degradation interaction), F(1,41) = 7.34, p = .01, but not with time-of-test (for 2 and 3-way interactions involving time-of-test group, Fs < 1.81).

Next, we investigated whether this long-lasting learning relies on the lateral orbitofrontal cortex (lOFC) as suggested by several prior studies using procedures that combine degradation with extinction^17–19^. We have previously argued^20^ that the lOFC supports contingency degradation through its role in representing internal states within cognitive maps of task space (i.e. the mental representation of the steps it takes to complete a task). We proposed that this role is particularly important when such states are required to parse competing contingency information^20,21^, as occurs during degradation learning. As illustrated in Figure 2 (top panel), this framework predicts that the lOFC divides original and degraded contingencies into separate states, such that inactivating it should impair degradation learning. This was tested in Experiment 2.

**Figure 2.**
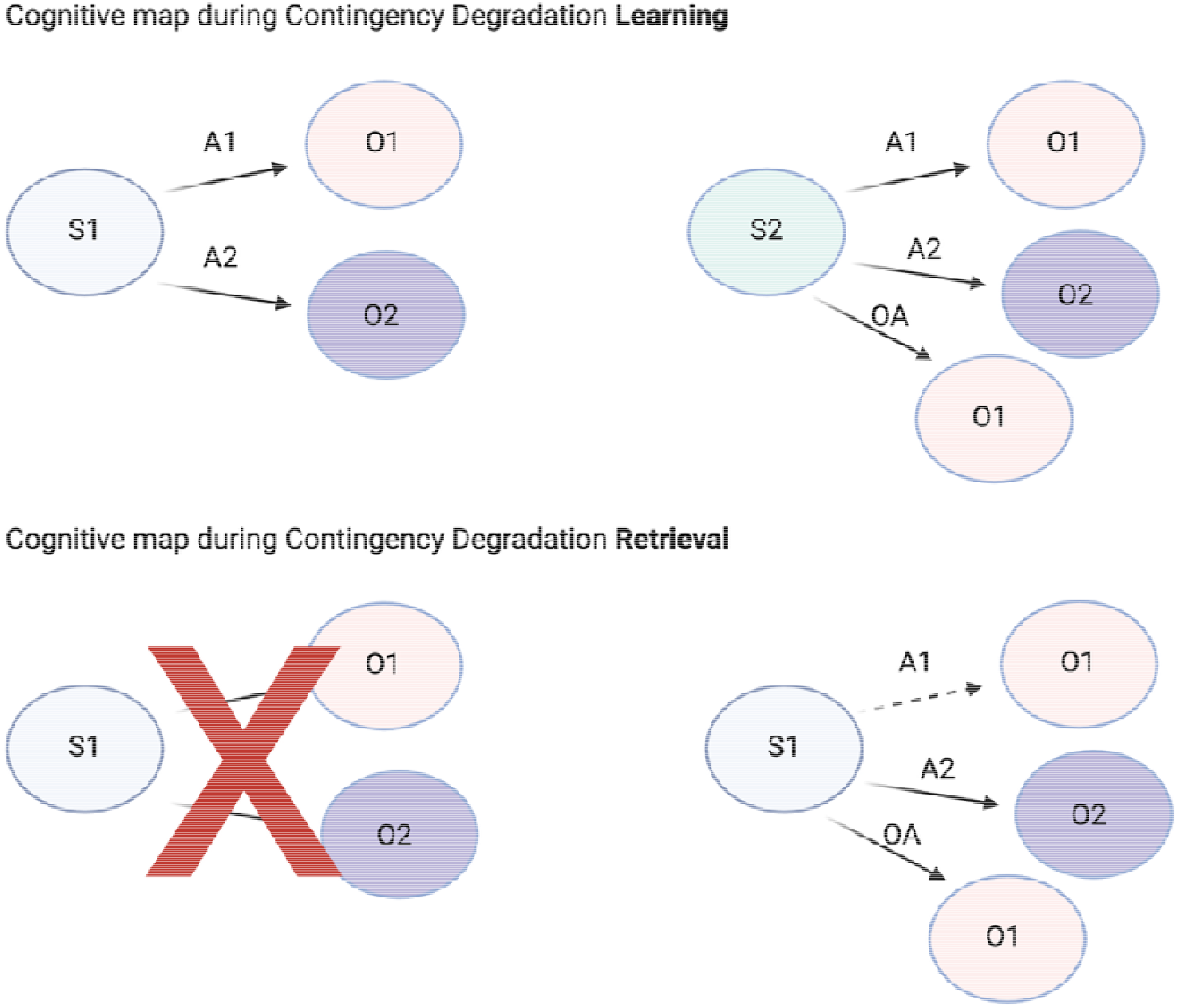
Proposed cognitive maps during contingency degradation learning and retrieval. Top panel: During initial lever press learning (top left), animals learn that in State 1 (S1), action 1 (A1) earns outcome 1 (O1) and action 2 (A2) earns outcome 2 (O2). During degradation learning (top right) animals learn that ‘other actions’ (OA; e.g. sniffing, exploring, withholding lever pressing) also produce O1, causing competition (see^22^ for a formal model of this process). It is proposed that the lOFC infers a new initial state (S2) to separate these contingencies from those learned in S1. Bottom panel: During retrieval, initial learning and its associated state have been overwritten (bottom left) and the A1-O1 contingency has been degraded (bottom right, denoted by a dashed arrow). As a result, separate state representations are no longer needed to resolve competition between initial and updated learning. Instead, retrieval of the updated contingencies can be guided entirely by observable cues, such as placement in the operant chamber, a process that does not rely on lOFC. Therefore, this account predicts that degradation learning but not retrieval relies on an intact lOFC.

Lateral OFC inactivation was achieved through the local chemogenetic transfection of glutamatergic neurons with inhibitory hM4Di designer receptors exclusively activated by designer drugs (DREADDs) paired with intraperitoneal injections of the ‘designer drug’ deschloroclozapine (DCZ, see supplemental for details). DREADDs transfection for Experiment 2 is shown in Figure 3A, and diagrammatic representations of transfection for each group are shown in Figures 3B (eGFP) and 3C (hM4Di). Both groups acquired initial lever pressing (Fig. 3D), however only the control group (eGFP+DCZ) went on to show evidence of degradation learning (Fig. 3E-F). During lever press acquisition there was a main effect of day F(1.276,19.143) = 25.53, p < .001, no group main effects or interactions, Fs < 1.148). During degradation, there were no main effects of degradation, F(1,15) = 1.2, p = .291, or group, F(1,15) = 1.6, p = 0.226, but there was a group x degradation interaction, F(1,15) = 4.86, p = 0.044, consisting of a significant simple effect in group eGFP+DCZ (NonDegraded > Degraded), F(1,15) = 5.136, p = .039, and not in group hM4Di+DCZ (NonDegraded = Degraded), F < 1. This result shows that an intact lOFC is required for degradation learning. Although performance during the learning phase was the focus for this experiment, we also tested rats drug-free and the general patterns of responding at test supported a lack of learning in group hM4Di+DCZ (See Supplemental Figure 1).

**Figure 3.**
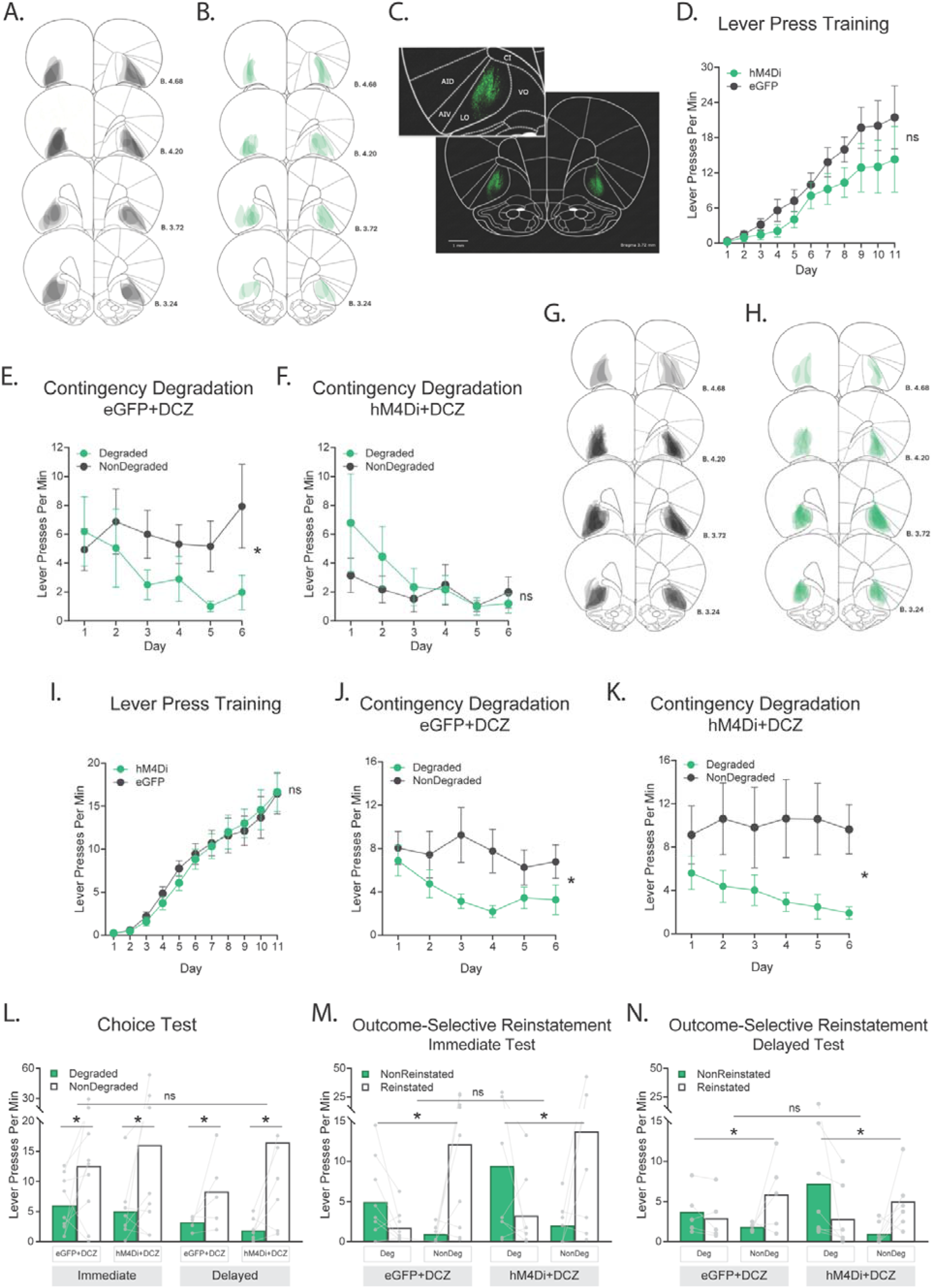
Lateral orbitofrontal cortex (lOFC) inactivation impairs contingency degradation learning but not retrieval. (A-F) Experiment 2 (role of lOFC in degradation learning). A) Verification of lOFC viral expression via fluorescence imaging. B-C) Overlapping viral placements for eGFP+DCZ (grey) and hM4Di+DCZ (green) groups. D-F) Lever presses per min (± SEM) during lever press training (D) and contingency degradation for eGFP+DCZ (E) and hM4Di+DCZ (F) groups. (G-M) Experiment 3 (role of lOFC in degradation retrieval). G-H) Overlapping viral placements for eGFP+DCZ (G) and hM4Di+DCZ (H) groups. I-K) Lever presses per min (± SEM) during lever press training (I) and contingency degradation for eGFP+DCZ (J) and hM4Di+DCZ (K) groups. L-M) Individual data points and mean lever presses during the choice test (L) and outcome-selective reinstatement (M-N). * p < .05.

In contrast to its role in degradation learning, we predicted that lOFC would not be required for the retrieval of degradation contingencies once learned (Fig. 2, bottom panel). This prediction follows the behavioural findings from Experiment 1, which suggest that by the end of degradation training, the initial action-outcome contingency has been forgotten, erased, or otherwise overwritten, such that it no longer competes with the degraded contingency for behavioural control at test. That is, unlike extinction where the original learning is retained and therefore spontaneously recovers and reinstates, after degradation, neither time nor outcome delivery was found to recover initial learning. In the absence of competition between initial and degraded contingencies, retrieval should no longer require lOFC-dependent state representations to separate them. This was tested in Experiment 3.

For this experiment, degradation learning occurred drug-free and was therefore intact in both groups (Fig. 3J-K). This was supported by a main effect of degradation, F(1,27) = 7.22, p = 0.012, and no group effects or interactions, Fs < 1. On the choice test, as predicted, lOFC inactivation had no effect on degradation retrieval at either time point, evidenced by intact degradation in all groups, main effect F(1,25) = 9.52, p = 0.005, and no group main effect or interactions, Fs <1 (Fig. 3L). Likewise, reinstatement replicated the pattern of responding observed in Experiment 1 in all groups, supported by a degradation × reinstatement interaction, F(1, 25) = 14.282, p < .001, with no group effects or interactions, Fs < 1 (Fig. 3M-N).

Altogether, these results reveal that instrumental contingency degradation reduces responding in a manner resistant to both spontaneous recovery and reinstatement, and that its learning but not retrieval depends on the lOFC. Thus, unlike extinction, degradation appears to overwrite the original action-outcome contingency. The dependence of degradation learning on lOFC observed in Experiment 2 nevertheless suggests that the original contingency is recalled during degradation learning, at least in the early stages, such that it competes with the degraded contingency and requires state segregation by lOFC. However, Experiment 3 indicates that this competition is no longer present at test. Determining precisely when original contingencies are overwritten during degradation learning therefore remains an important question for future studies.

Several additional questions remain. For instance, it will be important to identify how long degradation learning can persist for: will it survive a delay beyond the two weeks employed here? Is its persistence affected by longer (or shorter) periods of initial learning and/or degradation learning? Is a similar pattern of results observed in single lever-single outcome procedures, and in Pavlovian paradigms? Moreover, identifying whether lOFC involvement varies with any of these parameter changes and identifying where degradation learning is stored within the broader neural circuit remain critical goals, as present findings indicate only that it is stored somewhere other than the lOFC. These questions are important to answer if degradation is to be explored as a potential relapse-resistant treatment, particularly as individuals with compulsive disorders often display abnormal neural activity in their lOFC^23–25^ which could impair their ability to learn such treatments.

## Acknowledgements

We thank Melissa Sharpe for helpful feedback on this manuscript. We thank the technical staff at the technical staff at the Ernst Facility at the University of Technology Sydney. The authors acknowledge the use of the Nikon TI (Nikon corporation), Stellaris 8 (Leica Microsystems) and Axioscan Z1 slide scanning (Zeiss) microscopes in the Microbial Imaging Facility in the Faculty of Science at the University of Technology Sydney. We would like to thank Dr Amy Bottomley & A/Prof Louise Cole for their technical assistance. Figure 1a was created with BioRender.com.

## Funding

This work was supported by National Health and Medical Research Council grants GNT2003346 awarded to L.A.B, and GNT2028533 awarded to K.M.T. and L.A.B.

## Author contributions

M.M., M.D.K., and L.A.B., conceptualised and designed the research, M.M., and J.M.G performed the research (i.e. data curation), M.M., and L.A.B., analysed the data, L.A.B., and K.M.T., acquired the funding, M.M., A.C., J.M.G., M.D.K, K.M.T., and L.A.B. wrote the paper (original draft – M.M. and L.A.B., review and editing – A.C., J.M.G., M.D.K., and K.M.T.).

## Data availability statement

All the data, inputs, and outputs of statistical analysis for this manuscript are available on Open Science Framework at the following DOI: https://doi.org/10.17605/OSF.IO/HGZ7S.

## Competing Interests

None

## MATERIALS AND METHODS

### Experiment 1: Contingency degradation is resistant to spontaneous recovery and reinstatement

#### Animals

Experiment 1 used 43 Long–Evans rats (23 males and 20 females). Rats were 10 weeks or older at study onset and weighed 180–350 g. Animals were obtained from the Australian Research Centre (Perth, Australia) and housed in same-sex groups of three in transparent amber, ventilated plastic cages. The colony room was maintained at controlled temperature and humidity on a 12 h light/dark cycle, and all behavioural procedures were conducted during the light phase.

Before behavioural training commenced, rats were allowed one week to habituate to the laboratory environment with free access to food and water. Cages contained environmental enrichment, including a plastic tunnel, shredded paper, and a wooden gnawing object. During behavioural training and testing, food access was restricted to maintain animals at ∼85% of their free-feeding body weight, achieved by providing approx. 8-12g per female and 10-15g per male of maintenance diet per rat per day (note: this was highest at the start of lever press training to encourage exploration and pressing of the lever, but was increased the upper end of the range as instrumental responding became stable). All procedures were conducted in accordance with the Australian National Health and Medical Research Council (NHMRC) Code for the Care and Use of Animals for Scientific Purposes and were approved by the University of Technology Sydney Animal Ethics Committee (ETH21-6657).

#### Apparatus

All behavioural procedures took place in twelve identical sound-attenuating operant boxes (Med Associates, Inc.) located within individual sound-attenuating cubicles. The ceiling, back wall, and hinged front door of the operant chambers were made of a clear Plexiglas and the side wall were made of grey aluminium. Each chamber was equipped with a stainless-steel grid floor, as well as a recessed food magazine that was located at the base of one end wall, through which 20% sucrose-10% polycose solution (0.2 ml) and food pellets (45 mg; Bio-Serve, Frenchtown, NJ) could be delivered using a syringe pump and pellet dispenser, into separate compartments respectively. Two retractable levers could be inserted individually on the left and right sides of the magazine. An infrared light situated at the magazine opening was used to detect head entries. Constant illumination was provided by a 3-W, 24-V houselight situated at the top, centred on the left end wall opposite the magazine, and an electric fan fixed in the shell enclosure provided background noise (≈70 dB) throughout training and testing. The apparatus was controlled, and the data were recorded using Med-PC IV computer software (Med Associates, Inc.).

#### Behavioural Procedures

##### Magazine training

Magazine training was conducted on Day 1 to habituate rats to the operant chambers and to establish the food magazine as the site of outcome delivery. Rats were placed in the operant chambers, and the house light illuminated to signal the start of the session. No levers were extended during this session, but rats received 20 deliveries of food pellets and 20 deliveries of 20% sucrose solution on independent random-time 60 s (RT60) schedules. The house light was extinguished and the fan turned took approx. 25-30 mins.

##### Lever press acquisition

Lever-press training was conducted daily for 11 days (Days 2–12). Each session lasted a maximum of 50 minutes and comprised four 10-minute response periods: two on each lever, each separated by 2.5-minute time-out intervals during which levers were retracted and the house light turned off. Each 10-minute period terminated early if 20 outcomes were earned, such that rats could earn a maximum of 40 pellets and 40 sucrose overall per session.

Outcome assignments were counterbalanced across animals. For approximately half the rats, the left lever delivered pellets, and the right lever delivered a sucrose solution (a 20% sucrose plus 10% polycose solution), and the remaining rats received the opposite arrangement. For the first two days of lever press training, presses were reinforced on a continuous reinforcement schedule (CRF). Rats were then shifted to random ratio schedules: RR-5 for the next three days (reinforcement probability = 0.20), then to RR-10 for the following three days (probability = 0.10), and to RR-20 for the final three days (probability = 0.05).

##### Contingency degradation training

Contingency degradation training was conducted over the next 6 days (Days 13-18). Rats completed two 20-min sessions on each of these days, one on each lever, separated by a 1-2 hour interval, with the order of lever sessions alternated across days. During this phase, responses on each lever continued to produce their associated outcomes at the same programmed rate as during lever-press training (i.e. RR20: reinforcement probability = 0.05). For one lever, however, the corresponding outcome was also delivered non-contingently (“freely”), independent of responding, to degrade its contingency with the associated lever. The other lever was NonDegraded. To match response competition across Degraded and NonDegraded lever sessions, the Degraded outcome was delivered at the same rate in both (otherwise reduced responding on the Degraded lever could arise from increased magazine entries during Degraded sessinos rather than detection of the degradation contingency). The identity of the Degraded outcome (pellets vs sucrose) was counterbalanced across animals.

##### Choice test

The immediate group were tested one day after the last day of degradation training whereas the delay group were tested 2 weeks later. For these tests, rats were placed in the operant chamber with both levers extended but no outcomes were delivered. Again, the onset/offset of the houselight indicated the start and end of the session, respectively.

##### Outcome-selective reinstatement test

The day after the choice test, an outcome-selective reinstatement test was conducted. Rats first received 15 minutes of extinction in which both levers were extended no outcomes delivered. Next, there were four reinstatement trials, spaced four minutes apart. Each trial comprised a single, non-contingent delivery of either the sucrose solution or the grain pellet, following the same sequence for all rats: sucrose, pellet, pellet, sucrose. Pressing on each lever was measured during the two minutes before (Pre) and after (Post) each delivery.

##### Data and Statistical analysis

Behavioural data were collected automatically by Med-PC and exported to Microsoft Excel. Statistical analyses were conducted in IBM SPSS Statistics using the General Linear Model (GLM) framework with mixed-design (repeated-measures) ANOVA, with within- and between-subjects factors specified for each analysis. Where relevant, significant effects were followed up with planned comparisons using estimated marginal means (EMMs) and targeted pairwise comparisons aligned with a priori hypotheses. The alpha level was set at .05. Data are presented as mean ± standard error of the mean (SEM) and were averaged across counterbalanced conditions.

### Experiment 2: Learning contingency degradation depends on lateral orbitofrontal cortex

#### Animals

A total of 24 Long–Evans rats were used in Experiment 2 (12 males and 12 females). Following histological verification, three rats from the hM4Di+DCZ group and four rats from the eGFP+DCZ group were excluded due to injection misplacement. Final group sizes were therefore hM4Di+DCZ (n = 9) and eGFP+DCZ (n = 8). All housing and husbandry conditions were as described for Experiment 1.

#### Stereotaxic surgery

Rats were anaesthetised with inhalant isoflurane (5% for induction, 2–3% for maintenance) in oxygen delivered at a rate of 0.5 L/min throughout surgery. The rat’s head was shaved and then placed in a stereotaxic frame (Kopf), and 0.1 mL of the local anaesthetic bupivacaine administered at the incision site. An anterior–posterior incision was then made along the midline to expose the cranium, and the area surrounding bregma and lambda was cleaned and dried. Bilateral holes were drilled into the skull above the lOFC. Coordinates were anteroposterior (AP) = +3.7 mm from bregma, mediolateral (ML) = ±2.8 mm from the midline, and dorsoventral (DV) = −5.5 mm from the skull surface, according to the rat brain atlas (Paxinos & Watson, 2007). A 1.0 µL glass Hamilton syringe connected to an infusion pump (Pump 11 Elite Nanomite, Harvard Apparatus) was lowered to the target coordinates, and AAV was infused at a rate of 0.2 µL/min, delivering 1.0 µL per hemisphere. Specifically, rats in group hM4Di+DCZ were injected with pAAV-CaMKIIa-HA-hM4D(Gi)-IRES-eGFP (Addgene, item 50467-AAV2 at titer ≥ 2×10^12^ vg/mL) whereas rats in group eGFP+DCZ received pAAV5-CaMKIIa-eGFP (Addgene, item 50469-AAV2 at titer ≥ 7×10^12^ vg/mL), with assignments balanced within sex (i.e., half of the females and half of the males received each virus). The injector remained in place for 5 min following infusion to allow diffusion and minimise viral spread along the injector tract. During surgery, rats received a subcutaneous (s.c.) injection of 1mL/kg Meloxicam for analgesia and 3 mL warmed saline to compensate for fluid loss during surgery. An additional 1mL/kg Meloxicam (s.c.) was administered the following day to provide postoperative pain relief. Rats were allowed 7–10 days to recover in their home cages before commencing experimental procedures, during which time they were monitored and weighed daily.

##### Deschloroclozapine administration

DREADD agonist deschloroclozapine (DCZ) dihydrochloride (NIMH D-925) was acquired from National Institute of Mental Health (NIMH) through the NIMH Chemical Synthesis and Drug Supply Program. DCZ was diluted with sterile saline (SAL) (0.9% w/v NaCL) to a final injectable concentration of 0.1 mg/kg (at a volume of 1ml/kg) and was injected interperitoneally 20 min prior to contingency degradation training sessions. DCZ was always handled in dim/low light conditions (i.e. a single lamp in a darkened room) and freshly prepared on the morning of each test day.

##### Behavioural Procedures

Behavioural procedures were identical to those described for Experiment 1, except that all rats were tested immediately after degradation training (i.e. there were no delay tests).

##### Histological verification

Following the last day of behavioural procedures, rats were euthanised by CO_2_ inhalation and transcardially perfused with 4% paraformaldehyde (PFA) prepared in 0.1 M phosphate-buffered saline (PBS; pH 7.3–7.5). Brains were removed, post-fixed overnight in 4% PFA, and then placed in 30% sucrose for 3–5 days until they sank, indicating cryoprotection. Brains were then sectioned coronally at 40 μm through the lOFC, as defined by Paxinos and Watson (2014), using a cryostat (CM3050S; Leica Microsystems) maintained at approximately −20 °C. Sections were immediately immersed in cryoprotectant solution and stored at −20 °C. Later, three to four representative lOFC sections from each brain (Bregma +3.2 to +4.68) were collected and stained to verify the location and spread of transfection. During immunohistochemistry (IHC) sections were rinsed 3 times for 10 min in 0.1 M PBS, then submerged for 1 hr in PBS with 0.25% Triton and 3% BSA for blocking and permeabilisation. Sections were then placed in chicken anti-GFP (1:500, Millipore) as a primary antibody diluted in 0.25% Triton and 3% BSA in PBS for 24 hr at room temperature. Sections were then rinsed 3 times for 10 min in PBS and then placed in goat anti-chicken AF488 (Invitrogen) at 1:500 as a secondary antibody for 2 hr in PBS at room temperature. Sections were then rinsed three times with PBS for 10 min each and mounted from PBS using Vectashield mounting medium without DAPI (Vector Laboratories).

##### Imaging and immunofluorescence analysis

To determine AAV transfection and placement, a single image was taken of lOFC per hemisphere of each slice (6-10 images in total per brain region of each rat) on a Nikon TiE2 microscope using a 10x objective. Rats with misplaced, inconsistent, or lack of lOFC virus infusion were excluded from the study.

##### Data and statistical analysis

Data collection and statistical procedures were conducted as described for Experiment 1.

### Experiment 3: Contingency degradation retrieval does not depend on lateral orbitofrontal cortex

#### Animals

Experiment 3 used forty-one Long–Evans rats (20 male, 21 female). One rat died during surgery, two rats were determined to be statistical outliers (i.e. ± 2 SDs above the mean), and 9 animals with misplaced or insufficient AAV transfection were removed from the analysis. The resulting analysed sample comprised 15 rats in the hM4Di+DCZ group and 14 rats in the eGFP+DCZ group. After assigning rats to immediate and delayed groups there were four groups; (n=8, immediate (hM4Di+DCZ)), (n=9, immediate (eGFP+DCZ)), (n=7, Delayed hM4Di+DCZ)), (n=5, delayed (eGFP+DCZ)). All other housing and husbandry conditions were as described for Experiment 1.

#### Stereotaxic surgery and DCZ administration

All surgical and DCZ procedures were conducted identically to that described for Experiment 2, except that DCZ was administered 20 min prior to the choice and outcome selective reinstatement tests, whereas contingency degradation training occurred drug-free.

#### Behavioural procedures

Behavioural procedures were conducted identically to those described for Experiment 1.

#### Histological verification

Following the final behavioural session (the delayed outcome-selective reinstatement test), rats were perfused with 4% paraformaldehyde and their brains were collected. Tissue sectioning, immunohistochemistry, and microscopic verification of viral placement were conducted as described for Experiment 2.

#### Data and statistical analysis

Data collection and statistical procedures were conducted as described for Experiment 1.

**Supplemental Figure 1.**
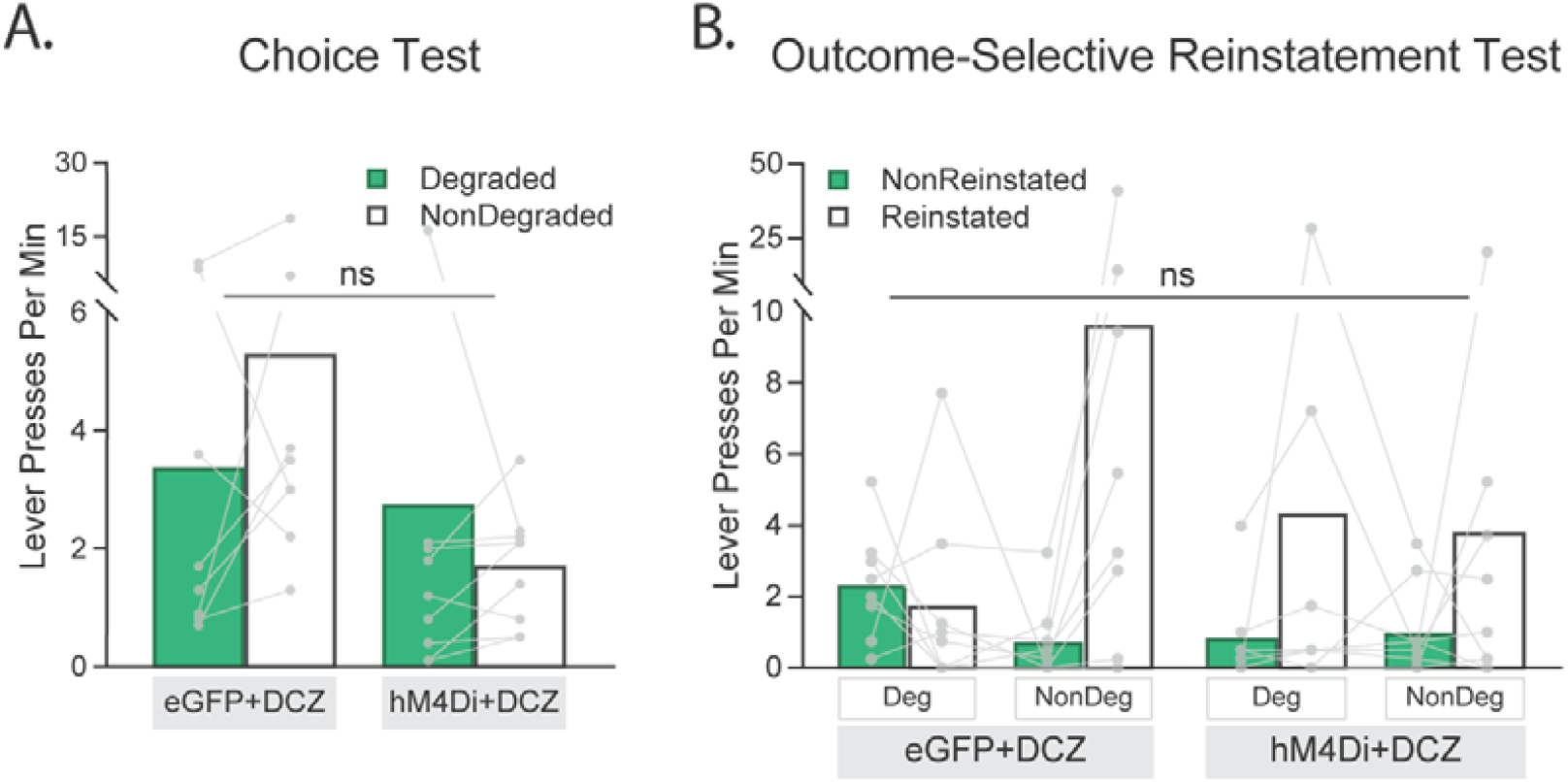
Test performance in Experiment 2 confirms a lack of learning in group hM4Di+DCZ. A) Individual data points showing performance during a 10 min choice test conducted in extinction. Group eGFP+DCZ show some evidence of a degradation effect (NonDegraded > Degraded) whereas group hM4Di+DCZ don’t (NonDegraded = Degraded), which prevented the observation of a degradation effect, F < 1. B) Individual data points showing performance during the outcome-selective reinstatement test. Group eGFP+DCZ show the same lack of reinstatement following delivery of the Degraded outcome (Reinstated = NonReinstated) as observed in Experiment 1, whereas group hM4Di+DCZ showed Reinstated > NonReinstated after delivery of both the Degraded and NonDegraded outcomes on the reinstatement – i.e. the expected pattern of responding had degradation not been learned. This is supported by a main effect of reinstatement, F(1,15) = 7.34, p = 0.016, but no group interactions, largest F for degradation x group x reinstatement F(1,15) = 2.22, p = 0.157.

## Notes

https://doi.org/10.17605/OSF.IO/HGZ7S

### Competing Interest Statement

The authors have declared no competing interest.

